# Enhanced hippocampal LTP but typical NMDA receptor and AMPA receptor function in a novel rat model of CDKL5 deficiency disorder

**DOI:** 10.1101/2022.06.29.497927

**Authors:** L Simões de Oliveira, HE O’Leary, S. Nawaz, R Loureiro, EC Davenport, P. Baxter, OR Dando, E. Perkins, SA Booker, GE Hardingham, MA Cousin, S Chattarji, TA Benke, DJA Wyllie, PC Kind

**Affiliations:** Centre for Discovery Brain Sciences, University of Edinburgh, Edinburgh, UK; Simons Initiative for the Developing Brain, Patrick Wild Centre, University of Edinburgh, UK; Centre for Brain Development and Repair, Instem, Bangalore, India; UK Dementia Research Institute, University of Edinburgh, UK; University of Colorado, School of Medicines, USA

## Abstract

Mutations in the X-linked gene cyclin-dependent kinase-like 5 (CDKL5) cause a severe neurological disorder characterised by early-onset epileptic seizures, autism and intellectual disability (ID). Impaired hippocampal function has been implicated in other models of monogenic forms of autism spectrum disorders and ID and is often linked to epilepsy and behavioural abnormalities. Many individuals with CDKL5 deficiency disorder (CDD) have null mutations and complete loss of CDKL5 protein, therefore in the current study we used a novel *Cdkl5* KO rat model to elucidate the impact of CDKL5 loss on cellular excitability and synaptic function of CA1 pyramidal cells (PCs). We hypothesised abnormal pre and/or post synaptic function underlie the enhanced LTP we observe in the hippocampus of *Cdkl5* KO rats. We tested this hypothesis using a combination of extracellular and whole-cell electrophysiological recordings, biochemistry, and histology. We show that NMDA receptor function and subunit expression are unaltered throughout development, and Ca^2+^ permeable AMPA receptor mediated currents are unchanged in *Cdkl5* KO rats. We observe reduced mEPSC frequency accompanied by increased spine density in basal dendrites of CA1 PCs, however we find no evidence supporting an increase in silent synapses when assessed using a minimal stimulation protocol in slices. Additionally, we found no change in paired-pulse ratio, consistent with normal release probability in *Cdkl5* KO rats and supported by typical expression of pre-synaptic proteins in synaptosome preparations. Together these data indicate a role for CDKL5 in hippocampal synaptic function and raise the possibility that altered intracellular signalling rather than synaptic deficits might contribute to the altered plasticity.

## Background

Mutations in the X-linked gene cyclin-dependent kinase-like 5 (CDKL5; MIM: 300203) cause a severe neurological disorder, estimated to affect 1 in 40,000 to 1 in 60,000 live births (Olson et al. 2019). Patients present with early onset seizures, sleep disturbances, motor impairments, autistic features and severe intellectual disability (ID) (Fehr et al. 2013; Olson et al. 2019).

Pathogenic mutations are predicted to result in loss of protein function and predominantly cluster in the catalytic domain of CDKL5 (Hector, Kalscheuer, et al. 2017), which is highly conserved across mice, rats and humans (Hector, Dando, et al. 2017; Hector et al. 2016). Identification of physiological substrates of CDKL5 has suggested a role in cytoskeleton organisation (Muñoz et al. 2018; Baltussen et al. 2018) which appears to be NMDA receptor dependent (Baltussen et al. 2018). In line with this role in cytoskeleton organisation, reduced dendritic complexity and altered spine distribution have been repeatedly reported in *Cdkl5* knock-out (KO) mice (Tang et al. 2017; Della Sala et al. 2016; Fuchs et al. 2014; Amendola et al. 2014). These anatomical phenotypes are frequently associated with altered synaptic function. Altered cellular and synaptic physiology has been reported in the hippocampus of a variety of mouse models of CDKL5 deficiency disorder (CDD). Altered long-term potentiation (LTP) has been observed in the hippocampus (Okuda et al. 2017) and cortex (Della Sala et al. 2016) of *Cdkl5* KO mice, with suggested mechanisms including an increase in calcium permeable AMPA receptors (Yennawar, White, and Jensen 2019) and NMDA receptor dysfunction (Tang et al. 2019; Okuda et al. 2017). These phenotypes are thought to underlie hippocampal-dependent learning and susceptibility to chemically induced seizures (Okuda et al. 2017, 2018; Tang et al. 2017).

NMDA receptor activation during development is known to influence synapse numbers and dendritic arborisation (Lüthi et al. 2001). Moreover, NMDA receptor subunit composition undergoes a developmental switch and has an important role in regulating AMPA receptor presence at synapses (Hall, Ripley, and Ghosh 2007). NMDA receptor development has not yet been studied in preclinical models of CDD. In fact, the role of CDKL5 in synaptic function during early postnatal development and juvenile stages is unknown, as most studies so far have focused on adult mice. Moreover, with conflicting reports from a variety of mouse models it is imperative to identify robust physiological phenotypes that cross the species barrier in order to identify disease mechanisms and therapeutic strategies which might translate to the human condition. In the current study, we report the generation of a novel *Cdkl5* KO rat model. We hypothesised that loss of CDKL5 leads to impaired synaptic function in the hippocampus of *Cdkl5* KO rats. We examined synaptic physiology and plasticity alongside cellular morphology using a combination of extracellular and whole cell electrophysiological recordings and histology. Whilst we found increased hippocampal LTP in *Cdkl5* KO rats, we found that NMDA receptors undergo a typical developmental trajectory and we do not observe an increase in calcium permeable AMPA receptor mediated currents that could contribute to the enhanced LTP. We observe reduced mEPSC frequency accompanied by increased spine density in basal dendrites of CA1 PCs, however we find no evidence supporting an increase in silent synapses when assessed using a minimal stimulation protocol in slices. Additionally, we found no change in paired-pulse ratio, consistent with normal release probability in *Cdkl5* KO rats. Overall our data, presents evidence supporting a role for CDKL5 in hippocampal synaptic function however the underlying mechanisms are still unclear and appear to be distinct to those previously reported in mouse models of CDD.

## Methods

### Breeding and animal husbandry: Edinburgh and Bangalore

All procedures were performed in line with the University of Edinburgh and Home Office guidelines under the 1986 Animals (Scientific Procedures) Act, CPCSEA (Government of India) and approved by the Animal Ethics Committee of the Institute for Stem Cell Science and Regenerative Medicine (inStem).

### Breeding and animal husbandry: Colorado

All studies conformed to the requirements of the National Institutes of Health *Guide for the Care and Use of Laboratory Rats* and were approved by the Institutional Animal Care and Use subcommittee of the University of Colorado Anschutz Medical Campus (protocol 00411). All rodents were housed in micro-isolator cages with water and chow available *ad libitum*.

Animals were bred in house on the Long Evans Hooded background and housed with littermates on a 12hr light/dark cycle with food and water *ad libitum*. Experiments were performed on wild-type (*Cdkl5*^*+/y*^) and *Cdkl5* KO (*Cdkl5*^*-/y*^) male rats at post-natal day (P) 28 to 35 unless otherwise stated. All experiments and data analyses were performed blind to genotype.

### *Cdkl5 KO rat generation and* genotyping

The CDKL5 KO rat model was created using CRISPR/Cas9 technology to introduce a 10 base pair (bp) deletion in exon 8 of the *Cdkl5* gene (Ensembl coordinates X:35674763-35674772, in the Rnor_6.0 genome assembly). An in-house PCR-based strategy was designed to genotype experimental animals produced from crossing Cdkl5 KO (*Cdkl5*^*-/y*^) males with wild-type females. Forward and reverse primers were generated flanking the bp deletion site in exon 8 of the rat CDKL5 gene (F1 and R), a third forward primer which anneals to the 10 bp deletion site in the WT allele (F2) and a further forward primer which anneals over the deleted 10 base pairs in the KO allele (F3) were also generated (Figure 1).

**Figure 1.**
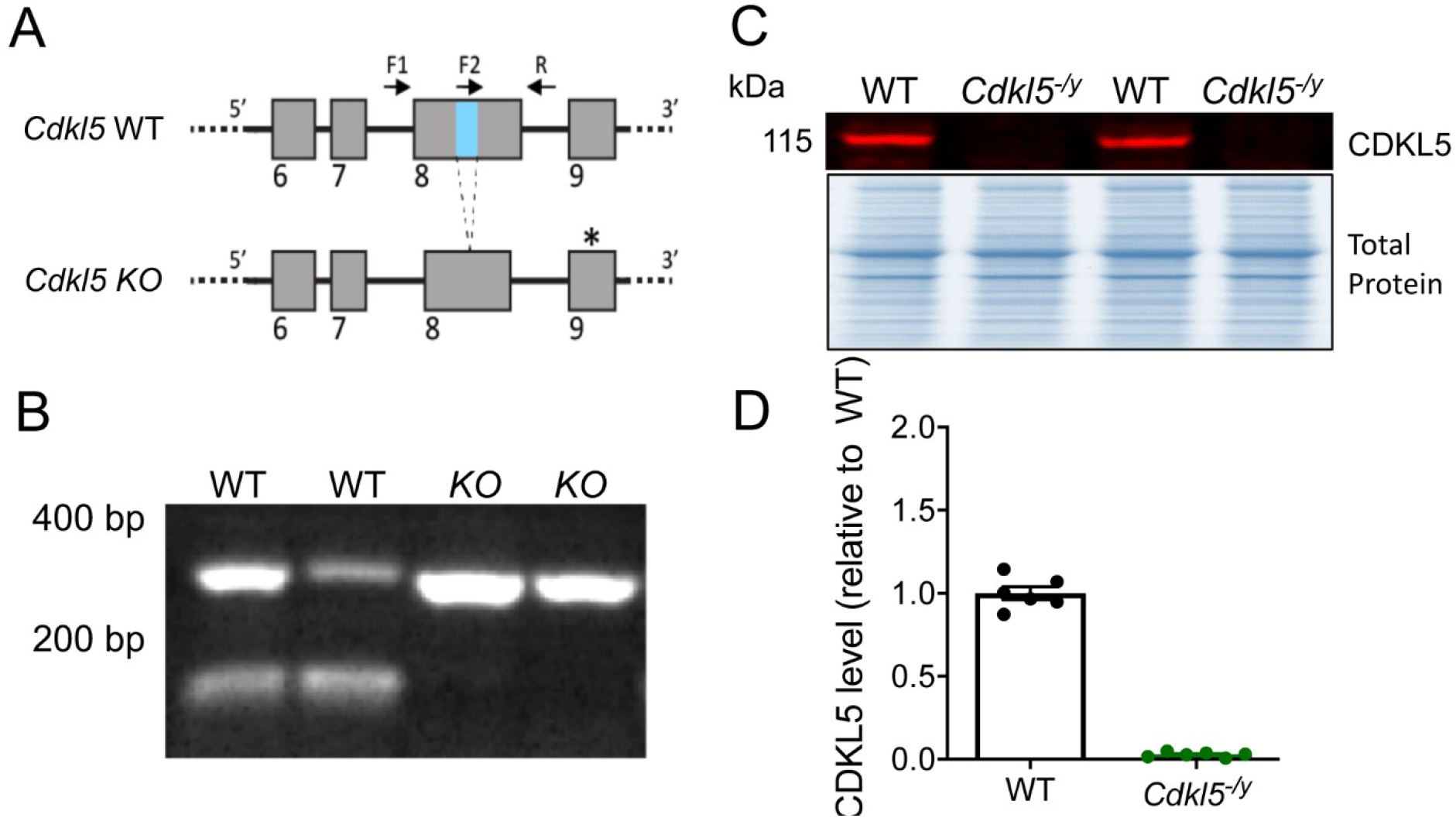
Validation of *Cdkl5* knock out rats. A - Schematic of the *Cdkl5* knockout strategy depicting the wild-type (WT) and null alleles. The null allele has a 10 base pair (bp) deletion in exon 8 (region shown in blue in WT allele), leading to a frame shift and an in frame, premature STOP codon forming in exon 9 (*). B - Genotyping results from male WT and Cdkl5 null animals. Higher band in WT and KO animals resulting from F1 and R primers product. Lower band in the WT samples resulting from F2 and R primer products is absent in the null samples due to the 10 bp deleted sequence. C - Western blot showing the absence of CDKL5 in hippocampal synaptosomes from the Cdkl5 null rats. D – Quantification of CDKL5 WB protein expression in hippocampal synaptosomes.

F1: 5’ -GGGCTTGTAGCAAATCCATCC- 3’

R: 5’ -AGCAAGCAGAGTTCTATTTTCCT- 3’

F2: 5’ -ATACGTGGCTACTCGGTGGTAC- 3’

F3: 5’ -CAGAATACGTGGCTACCGATC- 3’

To distinguish between DNA derived from wild-type and *Cdkl5*^*-/y*^ male littermates, primers F1, R and F2 were used in the same PCR reaction. Two bands were detected for wild-type male animals (356 and 135 bp) whereas only one band was detected for Cdkl5 KO male animals (346 bp) (Figure 1). To distinguish between DNA derived from wild-type and heterozygous female littermates, primers F1, R and F3 were used in the same reaction. One band was detected for wild-type female animals (356 bp) whereas two bands were detected for heterozygous female animals (356 and 129 bp).

Genomic DNA was extracted from fragments of tissue using the HotShot method. PCR was carried out as per the manufacturer’s guidelines for GoTaq G2 Polymerase (Promega, M784B) with an annealing temperature of 58°C and a 1 minute extension time.

Following initial validation experiments all genotyping was carried out by Transnetyx Inc.

### Acute slice preparation for electrophysiology

Acute brain slices were prepared from *Cdkl5*^*+/y*^ and *Cdkl5*^*-/y*^ at postnatal day (P) 28 to 35 (unless otherwise noted) similarly to previously described (Oliveira, Sumera, and Booker 2021). Briefly, rats were anesthetised with isofluorane and subsequently decapitated. The brain was rapidly removed and placed in ice-cold carbogenated (95 % O2/5 % CO2) sucrose-modified artificial cerebrospinal fluid (in mM: 87 NaCl, 2.5 KCl, 25 NaHCO3, 1.25 NaH2PO4, 25 glucose, 75 sucrose, 7 MgCl2, 0.5 CaCl2). 400 μm horizontal hippocampal slices were cut on a Vibratome (VT1200s, Leica, Germany). Slices recovered submerged in sucrose-ACSF at 34°C for 30 min and were then stored at room temperature until needed.

Alternatively, to assess NMDA receptor-mediated EPSCs throughout development and respective pharmacology, P7-22 *Cdkl5*^*-/y*^ and *Cdkl5*^*+/y*^ rats were rapidly decapitated and the brain removed. Parasagittal slices (400 μm) were prepared on a Leica VT 1200 microtome in ice cold solution containing (in mM) 206 Sucrose, 2.8 KCl, 1.25 NaH_2_PO_4_, 26 NaHCO_3_, 10 Glucose, 10 MgSO_4_, 2 NaAscorbate, 0.4 CaCl_2_, and 2.5 N-acetyl L-cysteine. Scalpel cuts were made to remove CA3 while retaining the CA1 region of the hippocampus with the overlying cortex and dentate gyrus intact for electrophysiology. Slices were then recovered > 60 min at room temperature in a submersion chamber in standard artificial Cerebral Spinal Fluid (aCSF), containing (in mM) 124 NaCl, 26 NaHCO_3_, 1.2 NaH_2_PO_4_, 10 D-glucose, 3 KCl, 2 NaAscorbate, 1 MgSO_4_, 2 CaCl_2_, and 2.5 N-acetyl L-cysteine) prior to all experiments. All solutions were oxygenated with 95% O_2_ - 5% CO_2_.

### Field LTP recordings

Slices were transferred to a submerged recording chamber perfused with warm carbogenated recording ACSF (in mM: 125 NaCl, 2.5 KCl, 25 NaHCO_3_, 1.25 NaH_2_PO_4_, 25 glucose, 1 MgCl_2_, 2 CaCl_2_) at a flow rate of 3-4 mL/min. Extracellular field recording electrode was filled with recording ACSF and placed in the *stratum radiatum* (*Str Rad*) of the CA1 region. Single pulses of electric stimulation (200 μs, 0.5 Hz) were delivered through a bipolar electrode (Ni:Cr) placed in the *Str Rad* to stimulate the Schaffer collateral (SC) pathway. Stimulus intensity was adjusted to produce 50% of the maximum field excitatory post-synaptic potential (fEPSP) amplitude. LTP was induced by tetanic stimulation (two trains of 1 s 100 Hz stimulation, 20 s inter-train interval, Komiyama et al., 2002) following 20 minutes of stable baseline. fEPSP slopes were normalised to baseline values and LTP magnitude reported as the average fEPSP slope in the final 10 min (50-60 min post-induction) of the recording divided by the average fEPSP slope during the baseline period. Data acquisition and analysis were performed on WinLTP (Anderson and Collingridge 2007).

### Whole-cell patch-clamp recordings

For whole-cell recordings, slices were transferred to a submerged recording chamber perfused with warm carbogenated recording ACSF (in mM: 125 NaCl, 2.5 KCl, 25 NaHCO_3_, 1.25 NaH_2_PO_4_, 25 glucose, 1 MgCl_2_, 2 CaCl_2_), at a flow rate of 6-8 mL/min. All recordings were performed at 31±1 ºC unless otherwise stated. Infrared differential inference contrast (IR-DIC) video microscopy, using a digital camera (Qimaging) mounted on an upright microscope (Olympus BX51WI) and a 40x (0.8NA) water immersion objective was used for all experiments. Recordings were obtained with a Multiclamp 700B (Molecular Devices) amplifier, signals were Bessel filtered online at 5 kHz and digitized at 20 kHz (Digidata1440, Molecular Devices) coupled to the Clampex software (pCLAMP™ Software, Molecular Devices). Recording pipettes were pulled from borosilicate glass capillaries (1.7 mm outer/1mm inner diameter, Harvard Apparatus, UK) on a horizontal electrode puller (P-97, Sutter Instruments, CA, USA), with resistance of 4-9 MΩ when filled with internal solution. For voltage-clamp recordings glass electrodes were filled with cesium based internal solution (in mM: 110 CsOH, 110 D-gluconic acid, 20 CsCl, 10 HEPES, 10 phospho-creatine, 4 MgATP, 4 NaCl, 0.3 Na_2_GTP, 0.2 EGTA, 5 QX314Cl) unless stated otherwise. A potassium gluconate based internal solution (in mM 120 K-gluconate, 20 KCl, 10 HEPES, 10 phospho-creatine, 4 MgATP, 4 NaCl, 0.3 Na_2_GTP, 2.7 biocytin, pH=7.4, Osm=290-310) was used for whole cell current clamp recordings.

Cells were rejected if series resistance >30 MΩ, or the series resistance changed by more than 20% over the course of the recording. No series resistance cancellation or junction potential corrections were performed.

### Evoked EPSCs

CA3 inputs to CA1 pyramidal cells were stimulated by placing a stimulating bipolar electrode (Ni:Cr or insulated tungsten) in the *Str rad* in hippocampal slices with the CA3 containing portion of the slice severed. A single 100 µs current pulse was delivered by an isolated constant current simulator (DS3, Digitimer.Ltd or WPI, Sarasota, FL). Evoked EPSCs were recorded in voltage-clamp using a cesium based intracellular solution.

#### NMDAR/AMPAR and paired-pulse ratio

AMPA receptor-mediated EPSCs were recorded at -70 mV in the presence of 50 μM picrotoxin to block GABAA receptors. The same cell was then held at +40 mV to record pharmacologically isolated NMDA receptor-mediated EPSCs in the presence of 50 μM picrotoxin and 10 μM CNQX. NMDAR/AMPAR ratios were calculated from peak amplitude of NMDA receptor and AMPA receptor-mediated EPSCs. We assessed paired-pulse ratio by evoking two EPSCs 50 ms apart, whilst holding the cell at -70 mV in the presence of 50 μM picrotoxin and calculating the ratio of the amplitude of the second EPSC relative to the first EPSC.

#### NMDA receptor development and pharmacology

A cesium based internal solution containing (in mM) 135 CsMeSO4, 10 HEPES, 10 BAPTA, 5 Qx314, 0.3 NaGTP, 4 Na_2_ATP, 4 MgCl_2_, and 0.1 spermine, pH 7.25 with 1 M CsOH was used. Recordings were performed at room temperature and extracellular solution was exchanged at a flow rate of 3-4 mL/min. AMPA receptor-mediated EPSCs were recorded at -70 mV, and NMDA receptor-mediated EPSCs were recorded at +40 mV. Peak current for NMDA receptor-mediated EPSCs was taken at 70 ms after the peak of the AMPA receptor-mediated EPSC. NMDA receptor sensitivity to block by GluN2B receptor antagonist Ro 25-6981 was determined by recording a 5 min baseline, followed by Ro 25-6981 (5 µM) perfusion onto the slice for 20 min.

#### AMPA-R I-V relationship

To assess the presence of calcium permeable AMPA receptors, AMPA receptor-mediated EPSCs were recorded in the presence of 50 μM picrotoxin to block GABAA receptors and 50 μM AP-5 to block NMDA receptors, over a range of voltages form -80 mV to +40 mV. Rectification index was calculated dividing peak EPSC amplitude at -60mV over peak EPSC amplitude at +40 mV. The same intracellular cesium based intracellular solution was used with added 0.1 mM spermine to maintain rectification of GluA2-lacking AMPA receptors.

#### Minimal stimulation

Minimal stimulation protocol was used to assess the presence of silent synapses. Once a reliable EPSC was identified at -70mV, stimulus amplitude was reduced until the synaptic response would fail in some of the trials, allowing for the stimulation of a single or a small number of synapses.

Following recording of 50 trials at a holding potential of -70 mV, corresponding to AMPA receptor-mediated EPSCs, the cell was depolarised to -40 mV, to reveal mixed AMPA and NMDA receptor-mediated EPSCs and an additional 50 trials were recorded. To determine response probability the traces for each holding potential were visually inspected and the number of traces with a visible EPSC was divided by the total number of traces for each cell. The ratio of response probability at the two holding potentials was used as an estimate for the relative abundance of silent synapses (Harlow et al. 2010; Isaac et al. 1997).

### Miniature EPSC recordings

Miniature EPSCs (mEPSCs) were recorded in voltage clamp while holding the cell at -70 mV, using a cesium gluconate based internal solution. Recordings were performed in recording ACSF in the presence of 50 μM picrotoxin and 300 nM TTX to block voltage gated sodium channels and consequently action potential firing. Analysis of mini EPSC frequency and amplitude over 1 minute of recording was performed using a template matching algorithm (Clements and Bekkers 1997) in Stimfit (Guzman, Schlögl, and Schmidt-Hieber 2014). A similar number of mini EPSC events was analysed for each condition.

### Intrinsic Physiology

Passive and active membrane properties were assessed to examine intrinsic excitability as previously described (Oliveira, Sumera, and Booker 2021). Passive membrane properties, including membrane time constant and input resistance, were measured from the voltage response to a 500 ms hyperpolarizing 10 pA step. Rheobase current and action potential (AP) firing frequency were determined from a series of depolarising current steps (0 to +400 pA, 500 ms) while holding the cell at -70mV with a bias current. AP properties were determined from the first AP elicited.

All analysis of electrophysiological data was performed using the open source software package Stimfit (Guzman, Schlögl, and Schmidt-Hieber 2014), blinded to genotype.

### Synaptosome preparation

*Cdkl5*^*+/y*^ and *Cdkl5*^*-/y*^ rats were killed by exposure to CO2 and decapitated. The hippocampus from each hemisphere was dissected in ice-cold 1x sucrose-EDTA buffer (0.32 M sucrose, 1 mM EDTA, 5 mM Tris, pH 7.4). The tissue was snap-frozen and stored at -80 °C until used for synaptosome preparation. On the day of preparation, the tissue was quickly thawed at 37 °C and homogenized in ice-cold 1x sucrose/EDTA buffer using 5-6 up-and-down strokes of a pre-chilled Teflon glass with motorized homogenizer (Dunkley, Jarvie, and Robinson 2008). Homogenates were centrifuged at 2800 rpm for 10 minutes at 4°C. The discontinuous (3% uppermost, 10% middle and 23% bottom) Percoll-density gradient was prepared prior to homogenization. The supernatant (S1) was added gently on 3% Percoll-sucrose (Percoll, P1644, Sigma Aldrich, UK) and centrifuged at 20,000 rpm for 8 min at 4°C. The fraction between 23% and 10% was collected and re-suspended in HEPES-Buffered-Krebs (HBK; in mM: 118.5 NaCl, 4.7 KCl, 1.18 MgSO_4_, 10 Glucose, 1 Na_2_HPO_4_, 20 HEPES, pH 7.4 balanced with Trizma) followed by centrifugation at 13,000 rpm for 15 min at 4°C. The pellet containing pure synaptosomes was dissolved in RIPA buffer (phosphatase inhibitor and protease inhibitor added). Protein quantification was performed with MicroBCA Assay kit (Pierce BCA protein estimation kit, 23225, ThermoFisher Scientific).

### Western Blots

Approximately 10 μg of synaptosome protein was separated on a precast gradient gel (NuPAGE 4-12% Bis-Tris Protein Gels, NP0336BOX, Thermo Fisher) and transferred to nitrocellulose membrane (AmershamTM Protran® Western Blotting Membrane, Nitrocellulose, GE10600002, Sigma Aldrich) using Bio-Rad transfer apparatus. Total proteins were stained with using reversible protein stain kit (Memcode 24580, Thermo Fisher Scientific) according to the manufacturer’s instructions. After removing the stain, membranes were blocked with 1:1 TBS1X: Odyssey Blocking Buffer (P/N-927-50003, LI-COR Biotech.) for an hour at room temperature, followed by overnight incubation with primary antibodies (CDKL5- 1:1000, #HPA002847, Atlas Antibodies-Sigma Aldrich; NMDAR1-1:1000, #700685, Thermo Fisher; NMDAR2A-1: 1000, #ab169873, Abcam; NMDAR2B- 1:1000, #610417, BD

Biosciences; PSD95- 1:2000, #76115, Abcam; GluR1- 1:1000, #MAB2263, Millipore; GluR2- 1:1000, #MABN1189, Millipore; RIM1/2- 1:2000, #140203, SYSY; Munc18-1- 1:2000, #116 011, SYSY; SNAP25- 1:1000, #111 011, SYSY; Syanpsin1- 1:1000, #ab64581, Abcam; Synaptophysin- 1:10,000, #ab32127, Abcam; VAMP2- 1: 10,000, #ab3347, Abcam) at 4°C. Membranes were washed with TBST1X (0.1% Tween 20), and incubated for an hour at room temperature with secondary antibodies (IRDye 800CW Goat anti Rabbit IgG- 1:10,000, #P/N 925-32211; IRDye 680LT Goat anti Mouse IgG- 1:10,000, #P/N 925-68020, LI-COR Biotechnology). Membranes were washed with TBST1X, dried and digitally scanned using Fc Odyssey Infrared Imaging System, LI-COR, UK Ltd. Odyssey software, Licor Image Studio Lite (LCOR Biosciences) was used to quantify individual bands. Data was normalised to respective total protein and then normalised to WT.

### Histology

Slices used for electrophysiology experiments were fixed in 4% paraformaldehyde (PFA) over night and stored in PBS (phosphate buffered saline) at 4 ºC until used for histology. Slices were washed in PBS and incubated in PBS with 0.3% triton-X and Alexa488 or Alexa568-conjugated streptavidin (1:500 dilution, Molecular probes, Invitrogen, USA) over night. Slices were then washed in PB and mounted on glass slides using Vectashield Hardset mounting medium (H-1400, Vector Labs).

### Image acquisition and analysis

To reconstruct cells and examine their morphology, multiple Z-stacks were taken in order to capture the entire biocytin-filled cell on an inverted confocal microscope (Axiovert LSM510, Zeiss) under a 20x Plan Neofluar (NA 0.5) objective (Zeiss). The Z-stacks obtained for a given cell were stitched using the 3D stitching plug in FIJI (ImageJ), and the cell was reconstructed using the Simple Neurite Tracer plug in (Longair, Baker, and Armstrong 2011). Sholl analysis was then performed on the skeletonised paths to examine dendritic complexity. To examine spine distribution in biocytin-filled cells, Z-stacks were taken from basal and apical (oblique and tuft) dendrites (2-3 dendrite sections per dendrite type per cell). Spines were imaged under a 63x Plan Apochromat (NA 1.4) oil immersion objective on an inverted confocal microscope (Axiovert LSM510, Zeiss), with a 2.8x zoom, 2x average line scan, 1024×1024 resolution, 0.14 μm Z step. Huygens Essential software (Scientific volume imaging, Netherlands) was used for deconvolution. The deconvolved images were used for analysis on FIJI (ImageJ). Z-projections of the deconvolved Z-stacks were used to manually count spines using the cell counter tool. For each dendrite section, the number of spines was normalised to the length of the section of dendrite analysed.

### Statistical analysis

All experiments and data analysis were performed blind to genotype. Where appropriate linear mixed effect models and general linear mixed effect models were implemented using the R package lme4 (Bates et al. 2014) on RStudio. Genotype was set as fixed effect and animal, slice (and cell where relevant) as random effects, allowing for direct measurement of genotype effect while accounting for the variability resulting from random effects. Where alternative statistical tests were used, Graphpad prism 7 was used to perform statistical comparisons across groups using two-tailed unpaired T-tests, repeated measures two-way ANOVA, or non-parametric tests as appropriate. In this case statistical testing was performed on animal averages to avoid pseudo-replication.

Details on sample size and statistical test used are presented in the results text and figure legends.

## Results

### Validation of CDKL5 KO rats

In collaboration with Horizon Discovery, the CDKL5 KO rat model was created using CRISPR/Cas9 technology to introduce a 10 base pair (bp) deletion in exon 8 of the *Cdkl5* gene (Ensembl coordinates X:35674763-35674772, in the Rnor_6.0 genome assembly). The deletion in constitutive exon 8 of the *Cdkl5* gene leads to a premature stop codon in constitutive exon 9 (Figure 1A). A genotyping strategy with primers flaking (F1, R) and overlapping (F2) the deletion site was used to distinguish WT and KO animals, with two bands detected for WT males (356 and 135 bp) whereas only one band was present for *Cdkl5*^*-/y*^ rats (346 bp) (Figure 1A-B). Examination of RNA-seq reads mapping to the *Cdkl5* locus confirmed the 10 bp deletion and revealed no cryptic splicing around the deletion. Thus, all transcripts produced from the locus are expected to contain the premature stop codon. The lack of CDKL5 protein expression was confirmed by western blot in hippocampal synaptosome preparations, where the 115 kDa band corresponding to CDKL5 is present in WT but not in *Cdkl5*^*-/y*^ rats (Figure 1C, quantified in 1D). Absence of CDKL5 protein in *Cdkl5*^*-/y*^ rats was further validated using proteomic analysis. *Cdkl5*^*-/y*^ rats were generally healthy with normal body weight and no overt behaviour phenotypes. *Cdkl5*^*-/y*^ rats did not exhibit observable spontaneous seizures.

### Enhanced hippocampal LTP in Cdkl5^-/y^ rats

To examine synaptic plasticity in the hippocampus of *Cdkl5*^*-/y*^ rats we performed extracellular field recordings in horizontal hippocampal slices from *Cdkl5*^*-/y*^ rats and their WT littermate controls aged P28 to P35. We measured the slope of the field EPSP evoked by stimulation of Schaffer collateral inputs to CA1 over a baseline period of 20 minutes and for 1 h following tetanic stimulation (Figure 2). Analysis of the LTP time course (Figure 2B) and the EPSP slope in the final 10 minutes of the recording as a percentage of the baseline EPSP slope (Figure 2C) revealed and enhanced LTP in *Cdkl5*^*-/y*^ rats (176.1 ± 5.6 %) relative to WT (138.3 ± 5.8 %, Two tailed T test, T=4.45, df=19, p=0.003). Interestingly, this is a transient effect as the magnitude of LTP is restored to WT levels by 12 weeks of age (Supplemental Figure 1).

**Figure 2.**
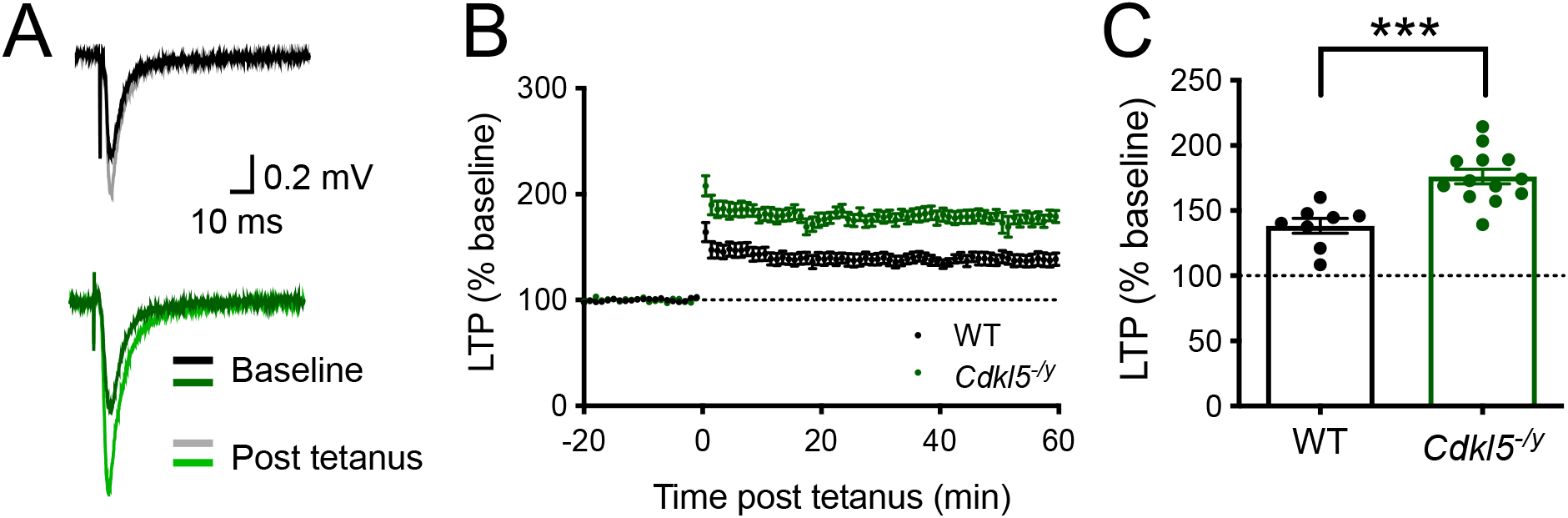
Hippocampal long term potentiation (LTP) in juvenile *Cdkl5*^*-/y*^ rats. **A -** Representative WT (upper) and *Cdkl5*^*-/y*^ (lower) fEPSP traces before (baseline) and after (post tetanus) LTP induction. **B -** Time-course showing long term potentiation (LTP) in the hippocampal CA1 induced by two trains with 100 pulses at 100Hz (20 seconds apart), resulting in a significant increase in LTP in *Cdkl5*^*-/y*^ rats when compared to WT. **C** – LTP in the final 10 minutes of the recording relative to baseline (WT n = 8 rats; *Cdkl5*^*-/y*^: n = 13 rats; *p<0.05 Two tailed T test, dots represent animal averages).

#### Unaltered NMDA receptor and AMPA receptpr function

Both AMPA receptor and NMDA receptor dysfunction and altered subunit composition have been implicated in abnormal LTP in mouse models of CDD (Okuda et al. 2017; Yennawar, White, and Jensen 2019). We assessed the NMDAR/AMPAR ratios of synaptic responses at the Schaffer collateral synapse of CA1, to test whether NMDA receptor function was altered in *Cdkl5*^*-/y*^ rats, possibly contributing to the enhanced LTP phenotype observed. AMPA receptor-mediated currents were recorded at a holding potential of -70 mV, whilst NMDA receptor-mediated EPSCs were recorded at +40 mV in the presence of the AMPA receptor antagonist CNQX (Figure 3A). CA1 pyramidal neurons in WT rats exhibited NMDAR/AMPAR ratio of 0.62 ± 0.08 (Figure 3B, B’) and an average NMDA receptor-mediated EPSC decay time of 85.69 ± 5.07 ms (Figure 3C, C’), which were unaltered in *Cdkl5-* ^*/y*^ rats (0.84 ± 0.14, GLMM: p=0.31; 105.3 ± 16.50 ms, p=0.78 Mann-Whitney test, respectively). These data indicate NMDA receptor function and subunit composition is unaltered in the absence of CDKL5. In line with the findings from electrophysiology experiments, the expression of NMDA receptor subunits GluN1, GluN2A and GluN2B was unaffected in the absence of CDKL5, as seen by comparable expression levels of these proteins in hippocampal synaptosome preparations from WT and *Cdkl5*^*-/y*^ rats (Figure 3 D-G).

**Figure 3.**
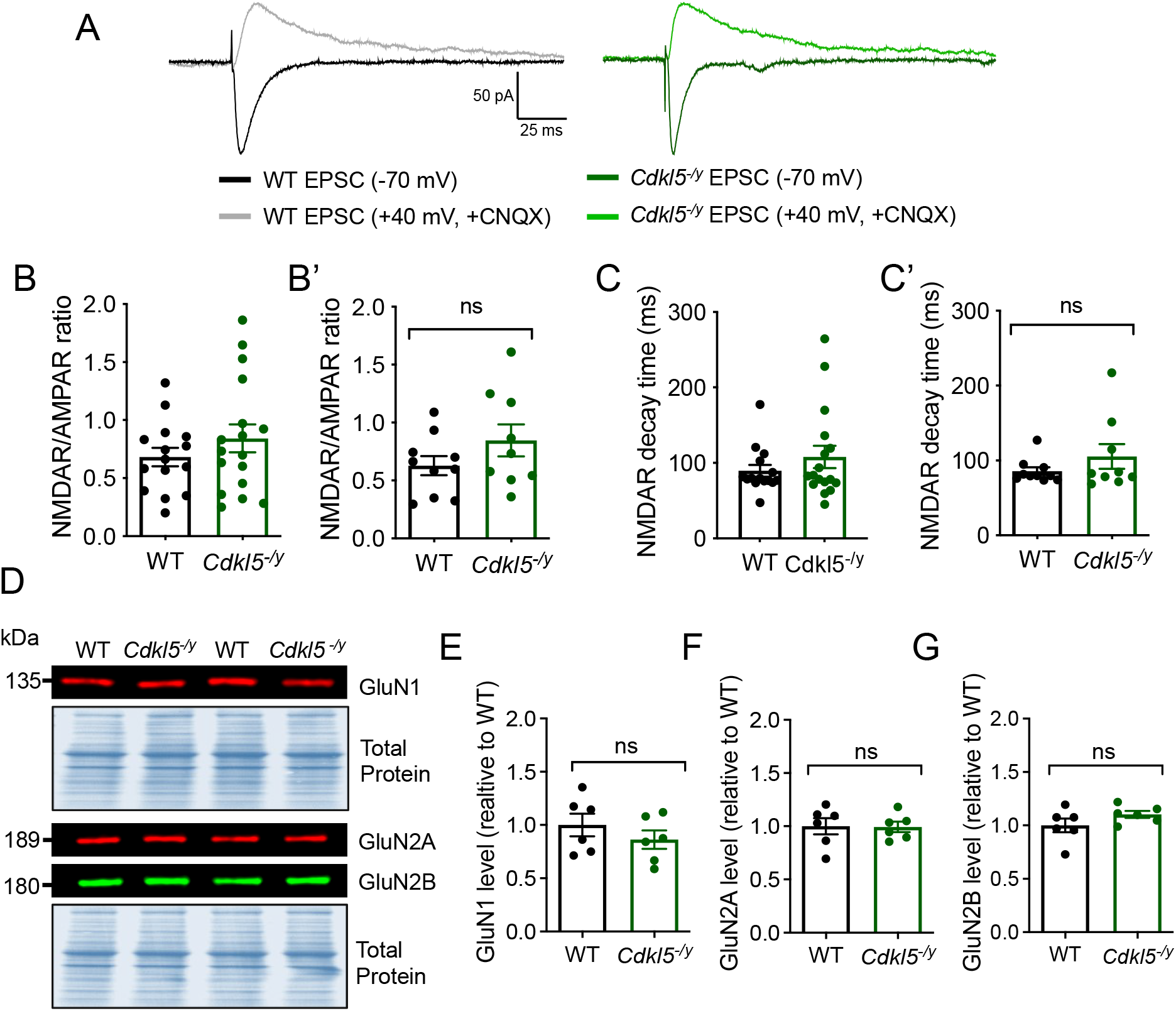
Unaltered NMDA receptor function and subunit composition in the hippocampus of P28-35 *Cdkl5*^*-/y*^ rats. **A –** Representative traces of AMPA receptor and NMDA receptor-mediated currents evoked by stimulating Schafer collateral inputs to CA1. **B, B**’ - NMDAR/AMPAR ratio (p=0.31 GLMM), **C, C’** – Pharmacologically isolated NMDA receptor-mediated EPSC decay time (p=0.78 Mann-Whitney U test performed on animal averages). Data shown as mean ± SEM (WT n = 10 rats / 14 cells; Cdkl5/y: n = 9 rats / 18 cells), dots represent individual cells (B, C) and respective animal averages (B’, C’). **D** – Representative western blot images from synaptosome preparations probed for NMDA receptor subunits GluN1, GluN2A, GluN2B and respective Total Protein stain. **E-G** – Quantification of protein expression level normalised to total protein and WT. Data shown as mean ± SEM. ns-p>0.05 Two-tailed T test.

Nonetheless, NMDA receptor subunit composition undergoes a developmental switch and has an important role in regulating AMPA receptor presence at synapses (Hall, Ripley, and Ghosh 2007). To the best of our knowledge, NMDA receptor development has not yet been studied in preclinical models of CDD, as studies conducted in mouse models have been largely restricted to adults. As such we examined NMDA receptor function over development to assess whether the developmental trajectory of NMDA receptor subunit composition is altered in *Cdkl5*^*-/y*^ rats, potentially contributing to long lasting effects at the circuit level.

NMDAR/AMPAR ratios and decay time constant of NMDA receptor-mediated EPSC were assessed in CA1 pyramidal cells from P7-22 rats (Figure 4A, B). In WT rats, the NMDAR/AMPAR ratio decreased from 1.24 ± 0.27 at P7-11 to 0.54 ± 0.13 at P18-22 (Figure 4A, 2-Way ANOVA age effect: F (2, 36) = 5.821, p=0.006), consistent with increased expression of AMPA receptor as development progresses (Pickard et al. 2000). This was accompanied by a reduction in decay time constant of the NMDA receptor-mediated EPSC from 562.9 ± 43.4 to 245.2 ± 20.3 over the same period (Figure 4B, 2-Way ANOVA age effect: F (2, 36) = 25.77, p<0.0001), consistent with an increased contribution of the NMDA receptor subunit GluN2A to synaptic transmission during development (Flint et al. 1997).

**Figure 4.**
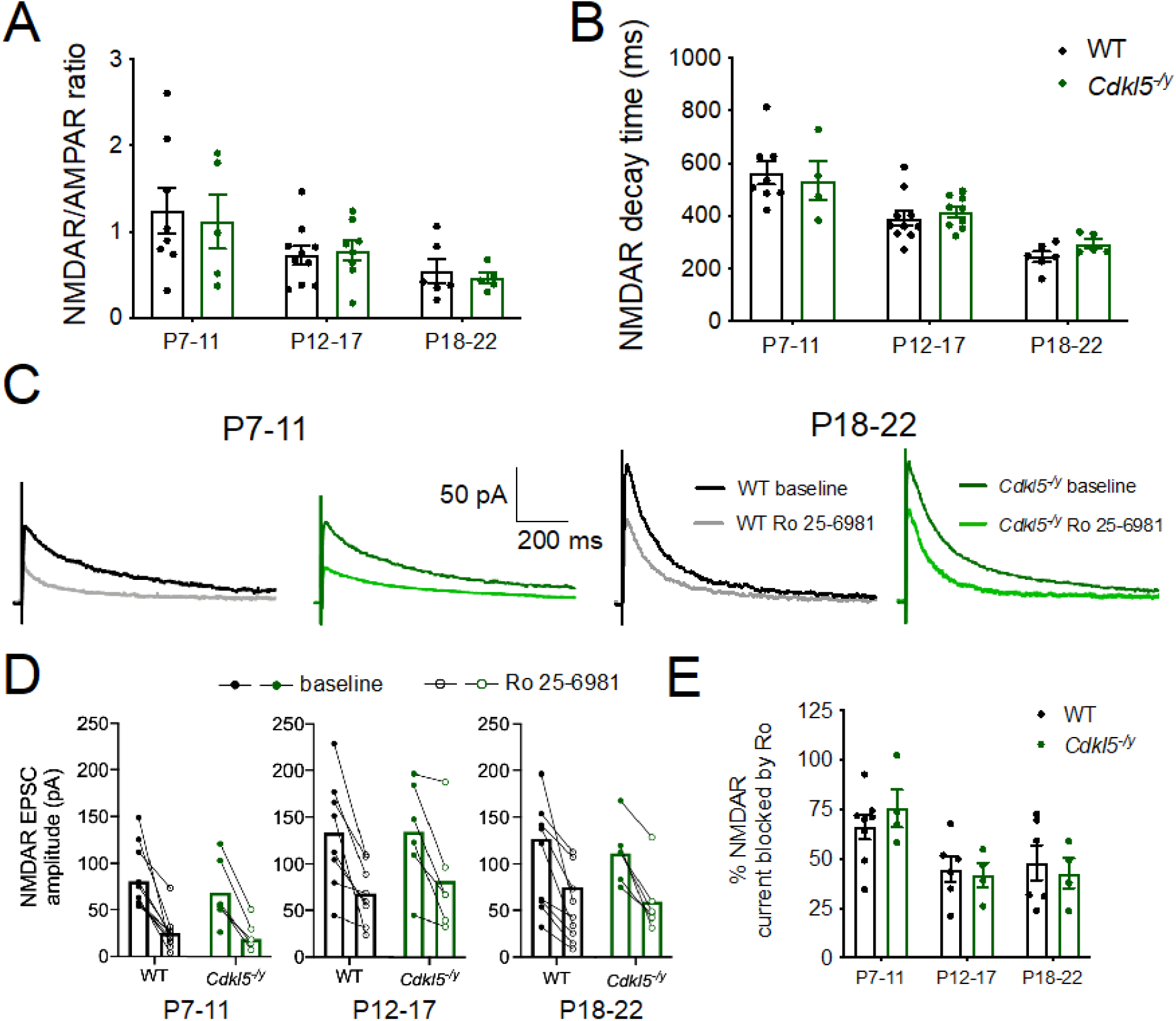
Typical NMDA receptor developmental trajectory in *Cdkl5*^*-/y*^ rats. **A –** NMDAR/AMPAR ratio in WT and Cdkl5^-/y^ rats aged P7 to P22. **B** - NMDAR decay time constant over development. **C** – Representative traces of NMDA receptor-mediated EPSCs in the presence or absence (baseline) of the GluN2B antagonist Ro 25-6981. **D** – NMDA receptor-mediated EPSC amplitude for individual cells before (full circles) and after (clear circles) Ro 25-6981 application, with recordings from each cell connected by a straight line across 3 age groups examined. **E** – Percentage of NMDA receptor current blocked by RO 25-6981 based on cells shown in D. All data shown as mean ± SEM, dots represent animal averages (except in D).

*Cdkl5*^*-/y*^ rats followed a similar developmental trajectory with NMDAR/AMPAR decreasing from 1.12 ± 0.32 to 0.46 ± 0.06, and decay time from 534.4 ± 73.12 ms to 294.0 ± 16.9 ms. This was not significantly different from WT when tested with a 2 Way ANOVA performed on animal averages (NMDAR/AMPAR - Interaction: F (2, 36) = 0.1397, p=0.87, genotype effect: F (1, 36) = 0.1043, p=0.75; decay time - interaction: F (2, 36) = 0.5142, p=0.60, genotype effect: F (1, 36) = 0.2326, p=0.63). Recordings in the presence of the GluN2B receptor antagonist Ro 25-6981 were performed to further examine the subunit composition of NMDA receptors during development (Figure 4 C-E). The percentage of NMDA receptor-mediated current blocked in the presence of RO 25-6981 decreased with age from 65.99 ± 6.18% at P7-11 to 48.06 ± 8.80% at P18-22 in WT rats. The block produced by Ro 25-6981 application was unaltered in *Cdkl5*^*-/y*^ rats (Figure 4E, Two-Way ANOVA Interaction: F (2, 26) = 0.542, p=0.59, genotype effect: F (1, 26) = 0.002, p=0.97). These data are further supported by the similar expression of NMDA receptor subunits in synaptosome preparations across genotypes in P14 rats (Supplemental Figure S2).

Overall, these data suggest that the developmental switch in NMDA receptor subunit composition and NMDA receptor contribution to synaptic transmission is unaltered in *Cdkl5*^*-/y*^ rats.

In addition to NMDA receptors, altered AMPA receptor subunit composition has been suggested as a potential mechanism underlying enhanced early-phase LTP in mouse models of CDD, where higher levels of calcium permeable (CP) GluA2-lacking AMPA receptors were observed (Yennawar, White, and Jensen 2019). To assess the relative abundance of CP-AMPA receptors, we recorded AMPA receptor-mediated EPSCs by stimulating the Schafer collateral inputs to CA1 and performing whole cell voltage clamp recordings from CA1 pyramidal cells in the presence of NMDA receptor and GABA-A receptor blockers (Figure 5). AMPA receptor-mediated EPSCs were recorded at a range of voltages from -80 mV to +40 mV in order to assess their current-voltage (I-V) relationship and rectification index. To maintain the intracellular polyamine block that confers inward rectification characteristic of CP-AMPA receptors, the intracellular solution in the recording pipette contained 0.1 mM of spermine (Kamboj, Swanson, and Cull-Candy 1995).

**Figure 5.**
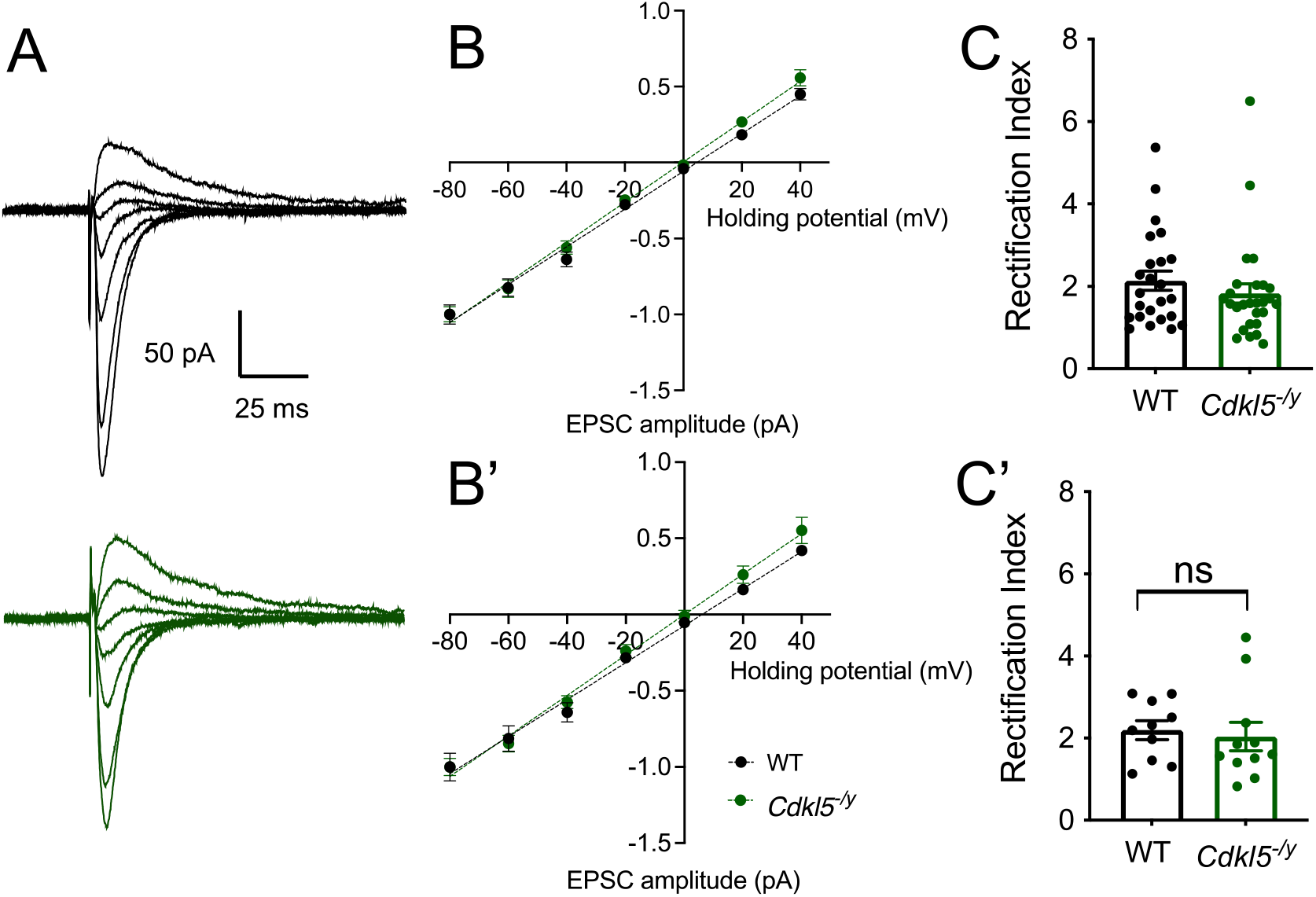
Unaltered AMPA receptor-mediated EPSC I-V relationship in CA1 pyramidal cells of Cdkl5^-/y^ rats. **A –** Representative traces of AMPA receptor-mediated currents from WT (upper, black) and *Cdkl5*^*-/y*^ (lower, green) recorded over a range of holding potentials (−80mV to +40 mV) in the presence of 0.1mM spermine in the intracellular solution. **B, B’ –** I-V relationship AMPA receptor-mediated EPSC normalised to EPSC amplitude at -80 mV holding potential (Genotype effect: F1,11=1.794, p = 0.21, Two-way ANOVA) **C –** Rectification index calculated as the ratio of EPSC amplitude at -60 mV over +40 mV. Data shown as mean ± SEM, data shown for individual cells (B, C) and animal averages (B’, C’) (WT n = 24 cells / 10 rats ; Cdkl5^-/y^: n = 27 cells / 11 rats)).

AMPA receptor-mediated EPSCs exhibited a linear current-voltage I-V relationship in WT neurons (Figure 5B, B’; linear regression: *y* = 0.0121*x* − 0.0729, r^2^=0.88), indicating no inward rectification and consistent with the high GluA2 expression in CA1 pyramidal cells (He et al., 1998, Pickard *et al*., 2000). In *Cdkl5*^*-/y*^ rats this linear relationship was maintained (linear regression: *y* = 0.0133*x* − 0.0002, r^2^=0.90) and did not differ to that observed in WT rats (Figure 5B, B’; F(1,143) = 2.3, p = 0.13, Sum-of-least squares F-test).

The rectification index calculated as the ratio of EPSC amplitude at the holding potentials of -60 mV and +40 mV was also unchanged in the absence of CDKL5 (WT: 2.19 ± 0.23, *Cdkl5*^*-/y*^: 2.04 ± 0.35, p=0.33 LMM). Together these data suggest that NMDA receptor and AMPA receptor mediated synaptic transmission are not affected in *Cdkl5*^*-/y*^ rats and therefore alterations to NMDA receptor and AMPA receptor are unlikely to contribute to the enhanced LTP observed in the rat model of CDD.

### Reduced mEPSC frequency and unaltered paired-pulse ratio in Cdkl5^-/y^ rats

In addition to post-synaptic mechanisms mediated by NMDA receptors and AMPA receptors, altered pre-synaptic function can contribute to abnormal synaptic transmission and altered synaptic plasticity. Indeed, CDKL5 has been implicated at the pre-synapse including through phosphorylation of the pre-synaptic protein amphiphysin-1 (Sekiguchi et al., 2013) and through its interaction with shootin1 which is thought to underlie normal axon specification (Nawaz et al. 2016). Moreover, reduced expression of the pre-synaptic marker synaptophysin has been reported in cellular models of CDD (Ricciardi et al. 2012; Fuchs et al. 2018). To assess further synaptic transmission in the absence of CDKL5, we recorded mEPSCs (Figure 6) and found a 30% reduction in mEPSC frequency from 5.43 ± 0.49 Hz in WT to 3.78 ± 0.34 Hz in *Cdkl5*^*-/y*^ rats (Figure 6 B-B’, LM p=0.02). This was accompanied by unaltered mEPSC amplitudes (Figure 6C-C’, WT: 11.51 ± 0.82 pA, *Cdkl5*^*-/y*^: 10.88 ± 0.69 pA, LMM p=0.64). As paired-pulse ratios (PPR) are commonly used to assess pre-synaptic release probability (Debanne et al. 1996), we next examined PPR of evoked EPSCs to determine whether the reduction in mEPSC frequency observed was a consequence of reduced pre-synaptic release probability. In WT rats, 2 pulses of electrical stimulation of Schafer collateral inputs to CA1 50 ms apart, resulted in a facilitating postsynaptic response with a PPR of 1.71 ± 0.11. In *Cdkl5*^*-/y*^ rats, EPSCs exhibited a PPR of 1.97 ± 0.18, unaltered relative to WT (Figure 6F-G’, LMM p=0.33).

**Figure 6.**
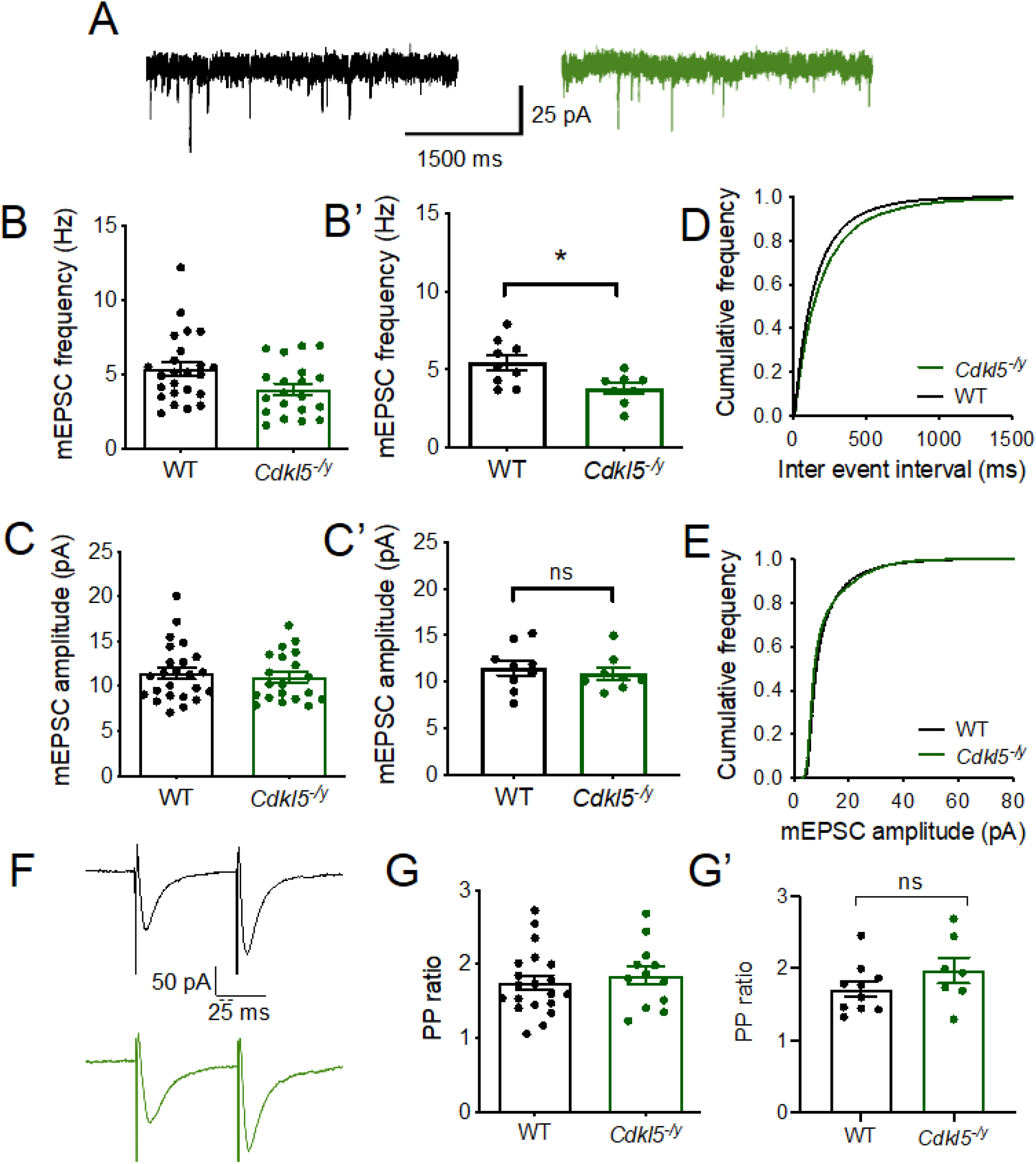
Reduced miniEPSC frequency and typical PPR in CA1 pyramidal cells from *Cdkl5*^*-/y*^ rats. **A -** Representative traces of mEPSC recordings from WT (left, black) and *Cdkl5*^*-/y*^ (right, green) rats. **B-B’** mEPSC frequency. **C-C’** mEPSC amplitude: WT n = 24 cells / 9 rats, *Cdkl5*^*-/y*^ n = 20 cells / 8 rats. **D** - Cumulative distribution of inter-event interval. **E** – Cumulative distribution of mEPSC amplitude. **F** - Representative traces of EPSCs evoked by PP stimulation of Schafer collateral inputs to CA1 pyramidal cells from WT (upper, black) and *Cdkl5*^*-/y*^ rats (lower, green). **G** - PPR of evoked (WT n= 20 cells / 10 rats, Cdkl5^-/y^ n= 10 cells / 7 rats) Data in bar charts shown as mean ± SEM (dots represent individual cells (B, C, G) or corresponding animal averages (B’, C’, D’).

### Typical dendritic morphology but increased spine density in basal dendrites of CA1 pyramidal cells

Altered dendritic morphology has previously been reported in mouse models of CDD across multiple brain areas (Okuda et al. 2018; Tang et al. 2017; Amendola et al. 2014). Moreover, dendritic morphology can have a profound impact on processing of synaptic inputs and consequently on circuit level function (Vetter, Roth, and Häusser 2001; Mainen and Sejnowski 1996). As such, we reconstructed biocytin-filled cells in order to examine dendritic arborisation and spine density in *Cdkl5-* ^*/y*^ rats (Figure 6, 7). In WT rats, biocytin-filled cells exhibited typical CA1 pyramidal cell morphology (Figure 7.A, (Amaral and Witter 1989; Bannister and Larkman 1995)). When examining the Sholl profile we found cell morphology to be unaltered in *Cdkl5*^*-/y*^ rats relative to WT controls (Figure 7.B, Two-way ANOVA on animal averages, Interaction: F 76,912 = 2.094, p<0.001, genotype effect: p=0.38). Total dendritic length (Figure 6C, C’, WT: 9882 ± 707 μm, *Cdkl5*^*-/y*^: 9455 ± 610 μm, Two-tailed T Test: T12=0.46, p=0.66) as well as total length of apical (Figure 7D, D’ WT: 6622 ± 520 μm, *Cdkl5*^*-/y*^: 5795 ± 519 μm, Two-tailed T Test: T12=1.12, p=0.28) and basal dendrites (Figure 7E, E’, WT: 3260 ± 257 μm, *Cdkl5*^*-/y*^: 3660 ± 275 μm, Two-tailed T Test: T12=1.06, p=0.31) were unchanged, indicating that overall dendritic complexity is not affected by the lack of CDKL5.

**Figure 7.**
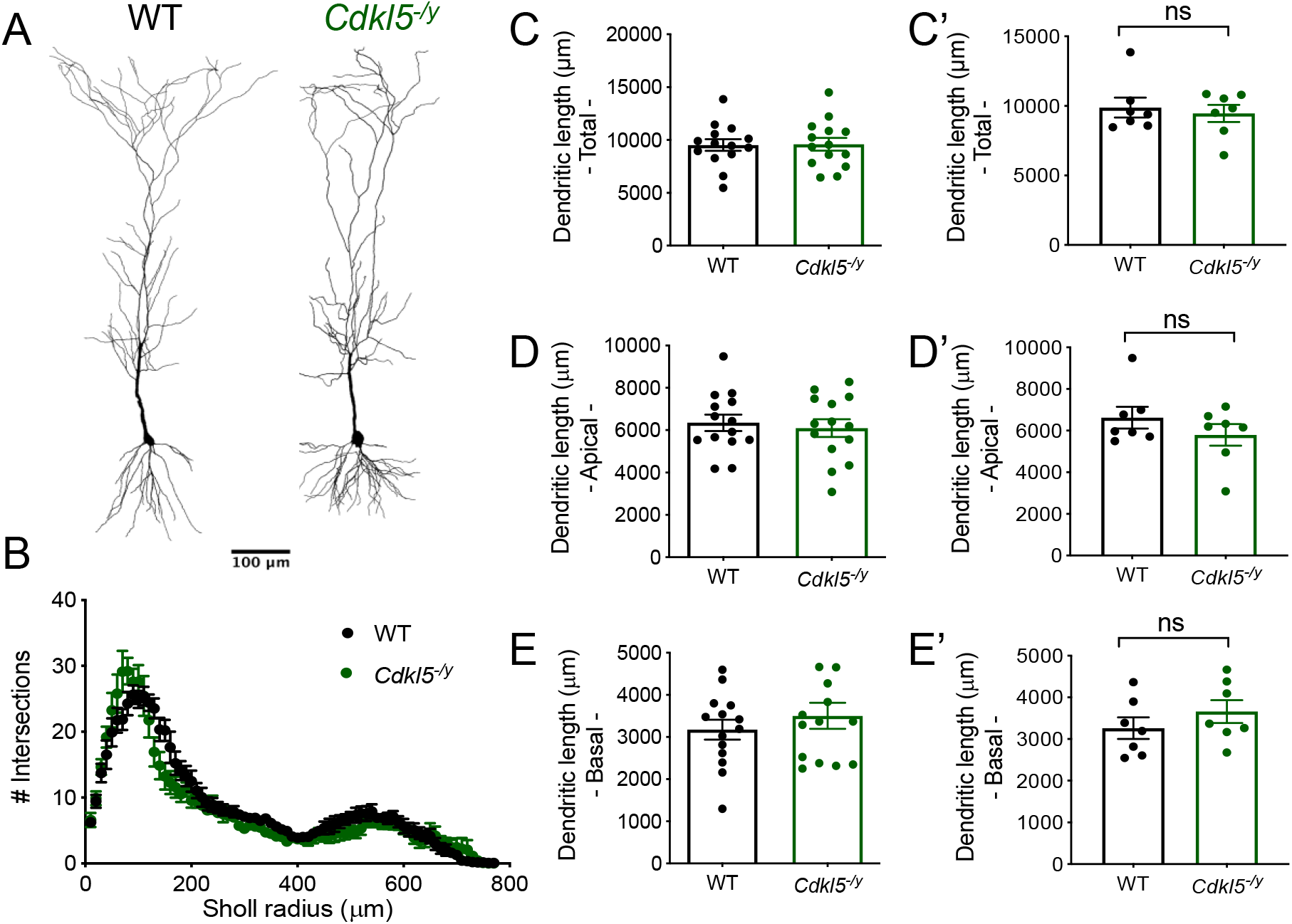
CA1 pyramidal cell morphology and spine density across multiple dendritic compartments. A – Example reconstruction of CA1 pyramidal cells from WT and Cdkl5^-/y^ rats filled with biocytin during whole cell patch clamp recordings. B – Sholl analysis of the dendritic arborisation (Two way ANOVA: Interaction: F 76,912 = 2.094, p<0.001, genotype effect p = 0.38). C – Total dendritic length, D – total length of basal dendrites, E – total length of apical dendrites. Data shown as mean ± SEM (WT - n=14 cells/7 rats, Cdkl5^-/y^ - n=14 cells/7 rats, dots represent animal averages, all p values > 0.05, Two tailed t-test)).

Despite no overall changes in gross dendritic morphology, we found a 19% increase in spine density in the basal dendrites of CA1 pyramidal cells from *Cdkl5*^*-/y*^ rats (Figure 8B-B’, 19.18 ± 1.18 spines/10 µm) relative to WT controls (16.08 ± 0.98 spines/10 µm, LMM p=0.04). Spine density did not differ between genotypes in the apical dendrites, oblique (Figure 8C-C’, WT: 20.02 ± 1.76 spines/10 µm, *Cdkl5*^*-/y*^: 18.30 ± 0.92 spines/10 µm, LMM, p =0.71) or tuft (Figure 8D-D’, WT: 8.71 ± 0.78 spines/10 µm, *Cdkl5-* ^*/y*^: 8.26 ± 0.82 spines/10 µm LMM p=0.71).

**Figure 8.**
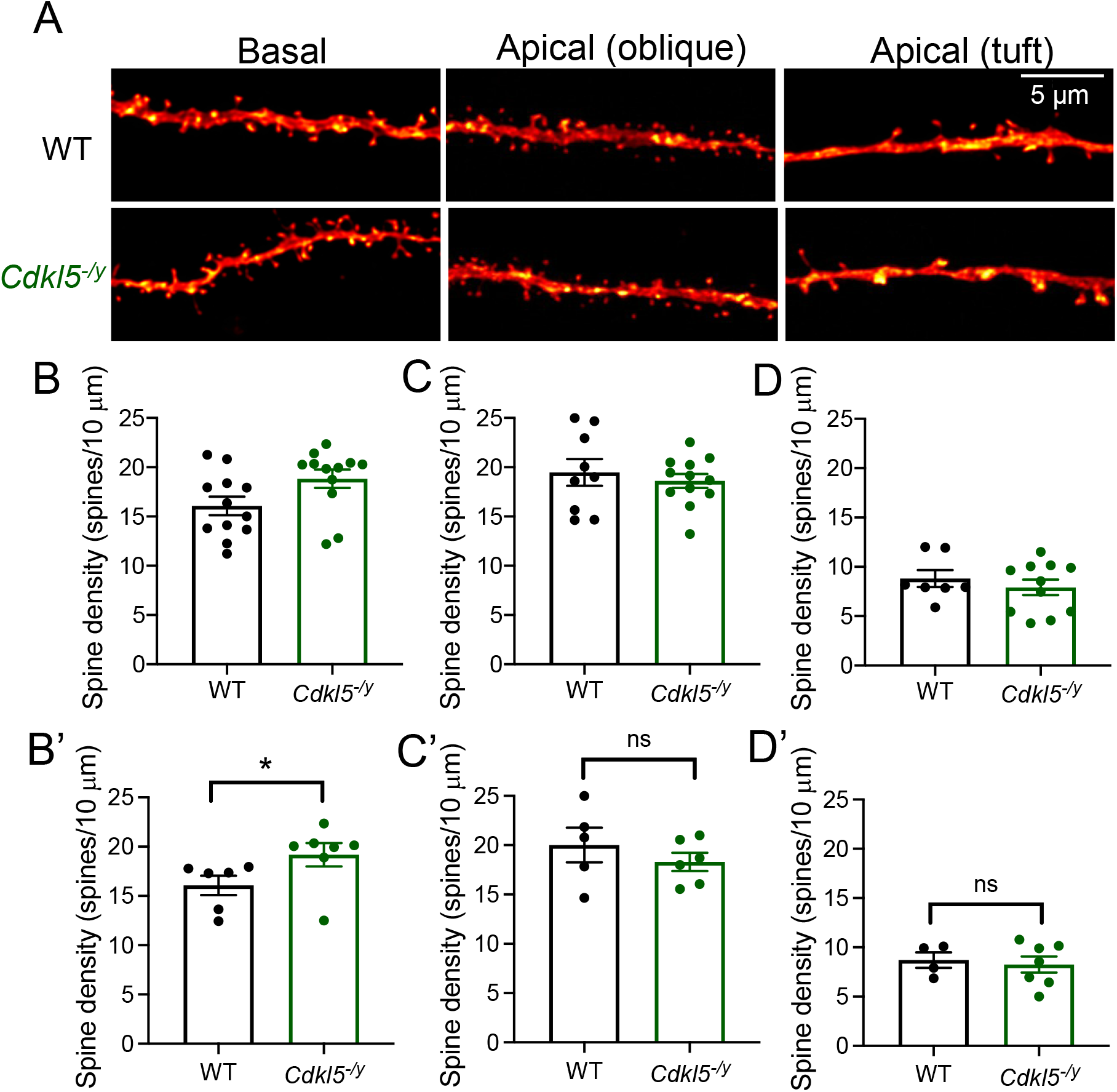
Spine density across dendritic compartments of CA1 pyramidal cells. A – Representative segments of basal and apical (oblique and tuft) dend*r*ites from CA1 pyramidal cells filled during whole*-*cell patch*-*clamp recordings. B – Spine density in basal dend*r*ites (WT: n = 12 cells/6 rats, *Cdkl5*^*-/y*^: n = 12 cells / 7 rats). C – Spine density in apical oblique dendrites (WT: n = 9 cells / 6 rats, *Cdkl5*^*-/y*^: n = 12 cells / 6 rats). D – Spine density in apical tuft dendrites (WT: n = 7 cells/4 rats, *Cdkl5*^*-/y*^: n = 11 cells / 7 rats). Data shown as mean ± SEM, dots represent cell (B, C, D) or animal averages (B’, C’, D’). *p<0.05, ns p>0.05 LMM.

### Unchanged relative abundance of silent synapses in Cdkl5^-/y^ rats

To determine whether the reduced mEPSC frequency and increased spine density observed resulted from an increase in the relative abundance of NMDA receptor-only silent synapses, we used minimal stimulation of Schaffer collateral inputs to activate a single or a small number or synapses onto CA1, thus resulting EPCSs or failures of synaptic transmission when recording AMPA receptor-mediated responses at a hyperpolarised holding potential (−70 mV). When the neuron is depolarised to +40 mV, mixed AMPA receptor and NMDA receptor-containing synapses as well as NMDA receptor-only containing synapses are activated (Figure 9A-B). Under these conditions, the ratio of response probability at +40 mV relative to -70 mV allows for an estimation of the relative abundance of silent synapses (Isaac et al. 1997). When recording at -70 mV, response probability was similar for WT and *Cdkl5*^*-/y*^ rats (WT: 0.66 ± 0.02, *Cdkl5*^*-/y*^ 0.73 ± 0.03, p=0.63). The response probability increased similarly in both genotypes when recording at +40 mV (WT: 0.79 ± 0.03, *Cdkl5*^*-/y*^ 0.89 ± 0.02, p=0.21 LMM), thus revealing the presence of silent synapses (Figure 9C). The similar ratio of response probability across genotypes (Figure 9D, WT: 1.23 ± 0.07, *Cdkl5*^*-/y*^ 1.21 ± 0.08, p=0.83 GLMM) is consistent with low levels of silent synapses in CA1 pyramidal cells (Racca et al. 2000) and indicates that the abundance of silent synapses is unaltered in the absence of CDKL5. Overall, these data suggest that altered abundance of silent synapses does not contribute to the LTP and mEPSC phenotypes observed in *Cdkl5*^*-/y*^ rats.

**Figure 9.**
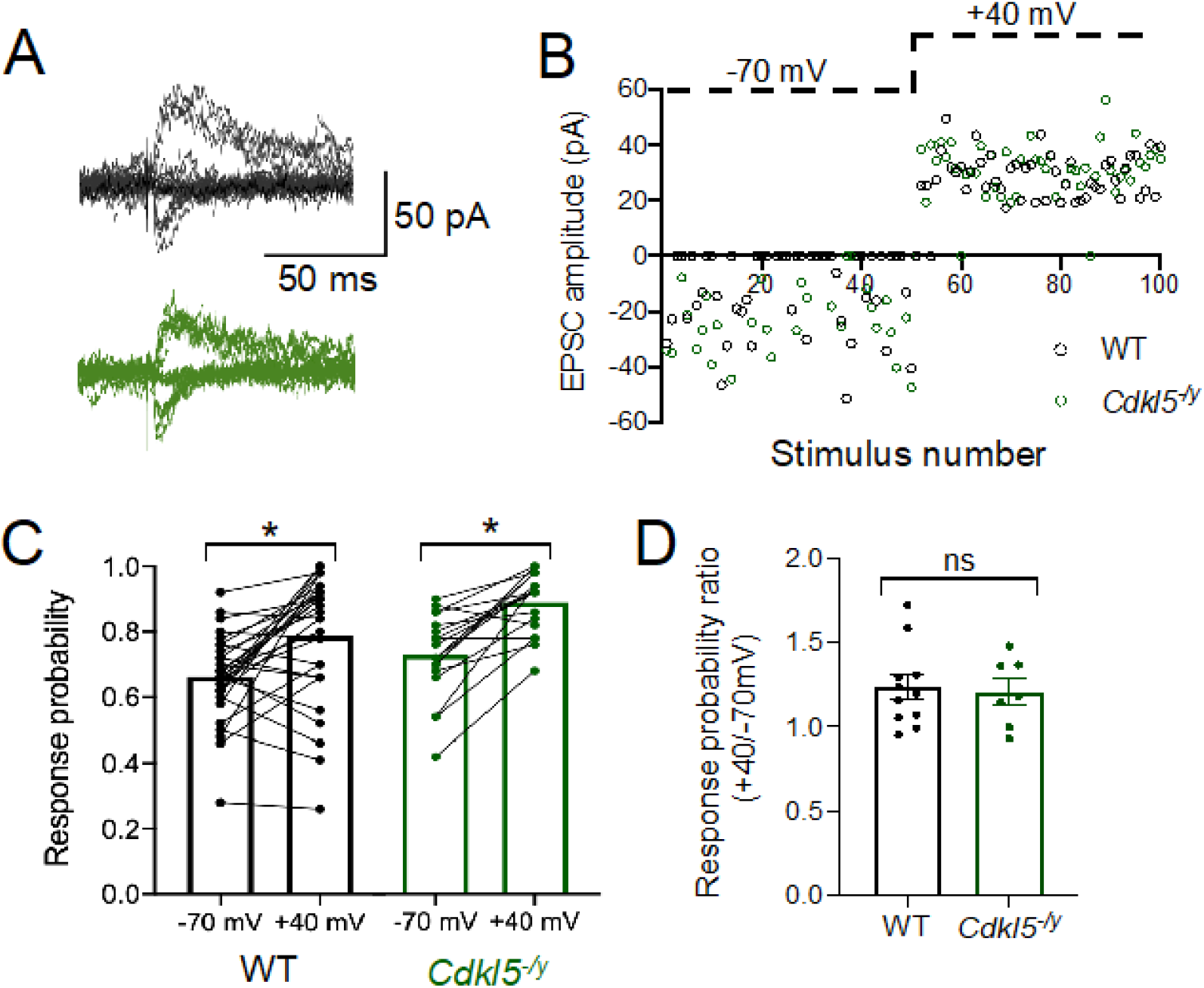
Minimal stimulation of CA3 inputs to CA1 pyramidal cells reveal no difference in silent synapses in *Cdkl5*^*-/y*^ rats. **A** - Representative traces of EPSCs recorded at -70 mV and +40 mV evoked by minimal stimulation of Schaffer collaterals. **B -** Example time-course of synaptic responses throughout a single WT and *Cdkl5*^*-/y*^ recording upon Schafer collateral stimulation. **C** - Response probability at -70 mV and +40 mV (data shown as cells, values for each cell connected by a black line. Two-way ANOVA Genotype effect: F_1,46_ = 5.16, p = 0.03, Holding potential effect: F_1,46_=38.50, p <0.0001). **D** - Ratio of the response probability at +40mV and -70 mV following minimal stimulation of Schaffer collaterals. (WT n= 30 cells / 11 rats, *Cdkl5*^*-/y*^ n=18 cells / 7 rats).

### CA1 pyramidal cells exhibit typical cellular excitability

In addition to synaptic transmission, we examined cellular excitability of CA1 pyramidal cells, as altered cellular excitability has been suggested to contribute to circuit level dysfunction in ASD/ID and epilepsy (Contractor, Klyachko, and Portera-Cailliau 2015; Clement et al. 2012).

CA1 pyramidal cells from WT rats exhibited a hyperpolarised resting membrane potential, fast membrane time constant and low input resistance (Table 1), in line with previous studies (Spruston and Johnston 1992; Staff et al. 2000). CA1 Pyramidal cells required 206 ± 20 pA of current injection to elicit the first AP (rheobase, Figure 10C-C’), and the number of APs fired increased with current injection thereafter until reaching a firing frequency of 22 ± 2 Hz in response to the maximum current injection step (400 pA, Figure 10A, D-D’). In *Cdkl5*^*-/y*^ rats, passive membrane properties (Table 1, Figure 10B-B’), rheobase current (LMM, p=0.91, Figure 10 C-C’) and overall AP firing in response to increasing current steps (Two-Way ANOVA F16,208=0.12, genotype effect: p=0.66, Figure 10D-D’), and were unaffected. These data indicate intrinsic neuronal excitability is unaffected in *Cdkl5*^*-/y*^ rats.

**Table 1.** Passive membrane properties of CA1 pyramidal cells are unaltered in *Cdkl5*^*-/y*^ rats.

| Physiological property | WT | $Cdkl5^{-/-}$ | p value (LMM) |
| --- | --- | --- | --- |
| Resting membrane potential (mV) | $-69.8 \pm 1.2$ | $-69.2 \pm 1.1$ | 0.56 |
| Input resistance (MΩ) | $68.1 \pm 6.8$ | $65.4 \pm 3.0$ | 0.64 |
| Membrane time constant (ms) | $19.1 \pm 1.0$ | $20.7 \pm 2.1$ | 0.64 |
| Capacitance (pF) | $299 \pm 23$ | $331 \pm 35$ | 0.69 |

**Figure 10.**
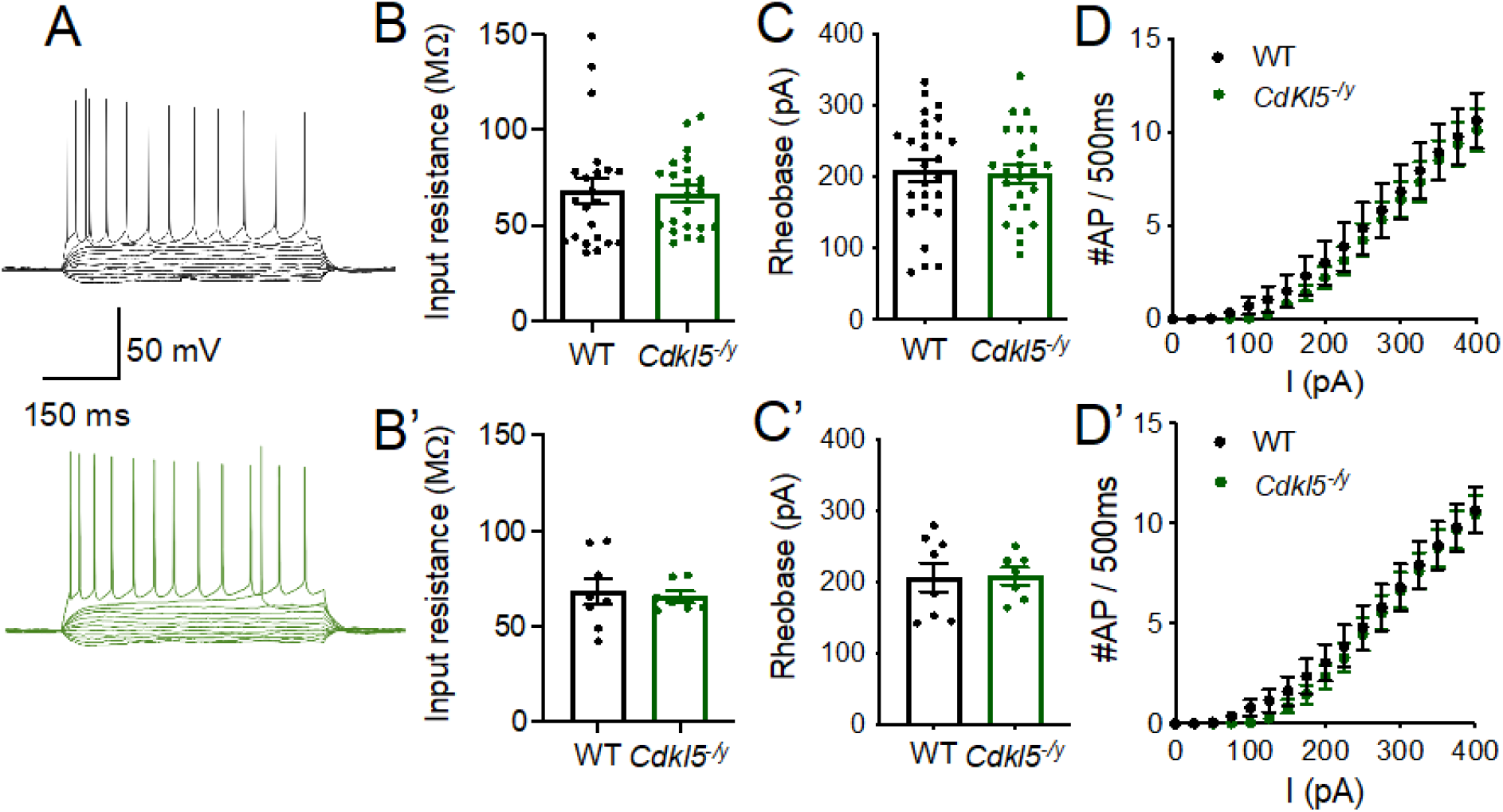
Typical excitability of CA1 pyramidal cells. **A** - Representative traces of whole cell recordings from WT (black, upper) and *Cdkl5*^*-/y*^ (green, lower) CA1 pyramidal cells in response to subsequent 25 pA steps. Traces shown from -100 pA to rheobase-1 and for the maximum firing frequency (I = 400 pA). **B, B’** – Input resistance. **C, C’** – rheobase current. **D, D’** - Action potential discharge in response to 500 ms long 25 pA current steps up to 400 pA (Two-way ANOVA genotype effect F16,208=0.12, p = 0.66). Data shown as mean ± SEM (WT – n = 26 cells/8 rats, Cdkl5/y – n = 24 cells/7 rats, dots represent single cells (B, D, D) or animal averages (B’, C’, D’).

## Discussion

In this study we described and validated a novel rat model of CDD, whereby targeting exon 8 of the *Cdkl5* gene resulted in complete absence of CDKL5 protein. Extracellular field recordings revealed enhanced LTP in the hippocampus of *Cdkl5*^*-/y*^ rats. However, extensive electrophysiological and biochemical characterization of NMDA receptors and AMPA receptors revealed no alteration in the functional properties of these receptors or the expression of their respective subunits. Further analysis of synaptic transmission in *Cdkl5*^*-/y*^ rats revealed a reduction in mEPSC frequency however, this finding was not accompanied by a change in PPR or an altered expression of hippocampal presynaptic proteins. Morphological characterisation of CA1 pyramidal cells with Sholl analysis revealed typical dendritic branching in *Cdkl5*^*-/y*^ rats. However, spine density was altered in a dendritic domain specific manner, with basal dendrites exhibiting higher spine density in *Cdkl5*^*-/y*^ rats relative to WT, while spine density was unchanged in apical dendrites. Despite this increase in spine density and reduced mEPSC frequency, minimal stimulation experiments revealed unaltered abundance of silent synapses in *Cdkl5-* ^*/y*^ rats. Cellular excitability was also largely unaffected in the absence of CDKL5.

*Mechanisms underlying enhanced hippocampal LTP are not conserved across mouse and rat models of CDD* In this study we found enhanced LTP in the hippocampus of *Cdkl5*^*-/y*^ rats, suggesting a role of CDKL5 in synaptic plasticity in this brain region. Whilst enhanced LTP has previously been reported in mouse models of CDD (Okuda et al. 2017; Yennawar, White, and Jensen 2019), the mechanisms previously suggested to contribute to this phenotype in mice are not translated to the rat model used in this study. In contrast to Okuda *et al*., 2017, we did not observe alteration in NMDAR/AMPAR ratio, NMDA receptor kinetics or subunit expression. Moreover, the developmental trajectory of NMDA receptor function and subunit expression can have long lasting impact on circuit level function and had not yet been characterised in rodent models of CDD. In this study we show NMDA receptor development to be unaffected in *Cdkl5*^*-/y*^ rats.

Furthermore, the contribution of GluA2 lacking AMPA receptors to synaptic transmission is also unaffected in *Cdkl5*^*-/y*^ rats contrary to what has been described in mouse models of CDD (Yennawar, White, and Jensen 2019). Indeed, the linear I-V relationship of AMPA receptor mediated EPSCs observed in WT and *Cdkl5*^*-/y*^ rats is consistent with the known high expression of GluA2 subunit in CA1 pyramidal cells (He et al., 1998), which confers low calcium permeability and no inward rectification (Jonas and Sakmann, 1992; Jonas et al., 1994).

In addition to species differences, other factors can contribute to the discrepancies observed between our findings and previous studies. Namely the ages of the animals tested and the nature of the genetic alteration leading to lack of CDKL5. In this study we focused on early post-natal development (P7 onwards) and juvenile ages (P28-35) due to the neurodevelopmental nature of CDD. However, the vast majority of studies conducted in pre-clinical models of CDD have focused on adult mice (*i*.*e*. older than 2 months, (Tang et al. 2017, 2019; Okuda et al. 2017; Amendola et al. 2014; Okuda et al. 2018; Wang et al. 2012)). Moreover, altered NMDA receptor function has been reported in constitutive knock out of *Cdkl5* achieved by targeting exon 2 (Okuda et al. 2017), calcium permeable AMPA receptors have been implicated in the R59X mutation knock in mouse model (Yennawar, White, and Jensen 2019), whilst the rat model used in this study results from targeting exon 8. Interestingly, discrepancies in behavioural phenotypes have been described across the variety of mouse models generated so far (Zhu and Xiong 2019).

Our findings suggest ionotropic glutamate receptor function and expression of synaptic proteins is intact in *Cdkl5*^*-/y*^ rats, therefore the cellular mechanisms underlying enhanced LTP in the rat model of CDD are yet to be understood. Work elucidating CDKL5 targets is still in its early stages, and there is no evidence of CDKL5 directly regulating signalling cascades downstream from LTP induction. However, downregulation of the mTOR signalling pathway has been reported across different mouse models of CDD (Schroeder et al., 2019; Amendola et al., 2014; Wang et al., 2012). Furthermore, altered mTOR signalling has been implicated in various other models of ASD/ID which also present with synaptic plasticity phenotypes (reviewed in Winden et al. (2018)), as such examination of this pathway might provide insight into a potential mechanism for the synaptic plasticity phenotype in *Cdkl5*^*-/y*^ rats.

### Excitatory synaptic transmission

In this study we report a reduction in mEPSC frequency in *Cdkl5*^*-/y*^ rats. mEPSCs are synaptic events resulting from the stochastic release of a single vesicle of neurotransmitter. Whilst mEPSC amplitude is a proxy for the number of receptors in the postsynaptic membrane, mEPSC frequency is a correlate for presynaptic release probability and/or synapse numbers. PPR is typically used to infer about presynaptic release probability (Debanne et al., 1996; Dobrunz et al., 1997), however the relationship between PPR and release probability is complex with studies showing that PPR can be maintained even when release probability is altered (Manita et al. 2007; Burke et al. 2018). In this study we find PPR to be unaltered at Schaffer Collateral synapses in CA1 in Cdkl5^-/y^ rats. Whilst this does not exclude the possibility of a presynaptic effect, together with unaltered expression levels of presynaptic proteins, these data suggest that release probability is unlikely to be affected in the hippocampus of Cdkl5^-/y^ rats.

As the reduction in mEPSC frequency was accompanied by an increase in spine density in basal dendrites, we used minimal stimulation to address the hypothesis that Cdkl5^-/y^ rats exhibit a greater abundance of silent synapses. Furthermore, altered abundance of silent synapses has previously been suggested to underlie abnormal synaptic plasticity in other models of ASD/ID with co-occurring epilepsy (Harlow et al. 2010). The response probability observed upon minimal stimulation of Schafer Collateral inputs to CA1 was consistent with the prevalence of silent synapses expected for CA1, based on anatomical studies (Racca et al., 2000), but no genotypic differences were found, indicating that the reduction in mEPSC frequency we observe cannot be explained by an increase in functionally silent synapses. Nonetheless, in this study we examined spine densities in biocytin filled cells as an estimate for number of excitatory synapses, however no synaptic markers were used to determine whether those spines are putative functional synapses. Therefore, it is plausible that despite an increase in spine density, *Cdkl5*^*-/y*^ rats do not exhibit an increased number of functional synapses but rather a reduction, thus explaining the reduction in mEPSC frequency observed. In fact, knock down of CDKL5 in neuronal cultures resulted in increased spine densities accompanied by a reduction in puncta of synaptic markers and reduced mEPSC frequency (Ricciardi et al. 2012).

#### Limitations

The study of CDD in rodents has been clouded by the lack of a seizure phenotype in the models developed so far. Whilst children with CDD present with early-onset epilepsy (Bahi-Buisson and Bienvenu 2012), this feature of CDD is not translated to rodent models of the disorder (Wang et al. 2012; Amendola et al. 2014; Okuda et al. 2017), including the rat model described in this study. Indeed spontaneous seizures have only been observed in aged heterozygous female mice (> 300 days), with the burden of epileptic spasms depending on the nature of the genetic alteration (Mulcahey et al. 2020). The lack of a seizure phenotype in ours and other models generated thus far casts doubts on whether rodent models can be useful in understanding the cellular and circuit level alterations underlying epilepsy in CDD. Nonetheless, CDKL5 protein function is still poorly understood and rodent models can provide useful tool to understanding the role of CDKL5 in neurons at the molecular and cellular level. The contribution of this study to the understanding of CDD is limited by the fact that only hemizygous male rats (where CDKL5 is completely absent) were examined, whilst most cases of CDD occur in heterozygous females. Clinically, the spectrum of severity is similar in males and females (Demarest et al., 2019; Siri et al., 2021; MacKay et al., 2021). However, studying CDKL5 function in heterozygous females is complicated by the random X chromosome inactivation leading to mosaicism. The lack of reliable antibodies to identify CDKL5 positive and negative cells and the lack of reporter lines where this can be done in real time pose a significant obstacle to determine the role of CDKL5 in neuronal function in heterozygous females. Therefore, the study of hemizygous males can provide a useful tool to understand CDKL5 function in a simplified system.

## Conclusion

This study described a novel rat model of CDD, of value to understand the role of CDKL5 in neurodevelopment in rodents. Moreover, the generation of this rat model provides valuable tool to the CDD research community. In combination with the existing mouse models, the *Cdkl5* KO rat can be used to identify robust cross species phenotypes that can be used as biomarkers when assessing potential therapeutics in preclinical models of CDD. This study provides evidence of a role of CDKL5 in excitatory synaptic transmission and synaptic plasticity in the hippocampus however the underlying mechanisms by which loss CDKL5 results in enhanced LTP and reduced mEPSC frequency remain to be elucidated.

